# Tractometry-Based Quantification of Along-Tract White-Matter Hemispheric Asymmetry in Alzheimer’s Disease

**DOI:** 10.64898/2026.01.23.701434

**Authors:** Bramsh Qamar Chandio, Yixue Feng, Iyad Ba Gari, Jonathan Davis Alibrando, Sophia I. Thomopoulos, Julio E. Villalón-Reina, Kenny Liou, Sunanda Somu, Hannah Yoo, Talia M. Nir, Eleftherios Garyfallidis, Eileen Lüders, Fang-Cheng Yeh, Neda Jahanshad, Paul M. Thompson

## Abstract

White-matter hemispheric asymmetry is a fundamental property of human brain organization and is known to change in aging, neurodevelopment, and neurodegenerative disorders. Tractometry analyzes diffusion-derived microstructural measures along the full length of tracts, localizing changes to specific tract-segments rather than collapsing tracts into a single value. Yet, existing frameworks lack a principled way to quantify left–right hemispheric asymmetries along homologous tracts. Here, we introduce an asymmetry-aware tractometry framework that integrates a symmetric white-matter atlas with BUAN (Bundle Analytics) to enable anatomically consistent, along-tract comparison of homologous pathways. By defining homologous bundles with a shared template and consistent orientation, each left-hemisphere segment is directly matched to its right-hemisphere counterpart, enabling principled, segment-wise comparison and revealing spatially localized asymmetries along-tract. Applying this framework to diffusion MRI data from the Alzheimer’s Disease Neuroimaging Initiative (ADNI) comprising 1,215 subjects, we demonstrate how this approach reveals systematic left–right asymmetries across major white-matter pathways and show how these patterns differentiate cognitively normal (CN) individuals from those with mild cognitive impairment (MCI) and dementia. This method provides a sensitive and anatomically grounded tool for studying hemispheric specialization and its disruption in aging and disease, and establishes a general approach for asymmetry-aware tractometry in population neuroimaging studies.

## I. Introduction

Hemispheric asymmetry is a defining feature of the human brain, underlying specialization in language, motor control, and cognition [1]–[5], to name a few. Structural differences between the left and right hemispheres emerge early in development and evolve across the lifespan [6], [7], and disruptions to these patterns have been linked to a wide range of neurological and neuropsychiatric conditions, including Alzheimer’s disease [8], Parkinson’s disease [9], and developmental disorders [10], [11]. While cortical asymmetries have been extensively studied, much less is known about asymmetry in the brain’s white-matter architecture, particularly along the full extent of individual fiber pathways.

Diffusion MRI (dMRI) and tractography provide a unique opportunity to study white-matter organization *in vivo* by reconstructing whole-brain fiber pathways and enabling tract-specific analyses. Tractometry methods [12]–[14] project microstructural measures, such as fractional anisotropy (FA) and various diffusivity indices, along the length of segmented bundles, yielding spatially resolved profiles that can be compared across subjects and groups. These approaches have proven powerful for detecting localized alterations in development [15], aging [16], and disease [17]. However, existing tractometry frameworks are not designed to explicitly model hemispheric symmetry. Left and right homologous bundles are typically analyzed independently, making it difficult to directly quantify spatially aligned asymmetries along the tract.

A principled analysis of white-matter asymmetry requires two key components: (i) a symmetric anatomical reference that enables homologous left–right correspondence, and (ii) a tractometry framework capable of capturing localized, along-tract differences. In this work, we introduce a new asymmetry-aware tractometry approach that integrates a symmetric white-matter atlas [18] with BUndle ANalytics (BUAN) tractometry [14]. By registering homologous left and right bundles into a common symmetric space, we establish segment correspondence along each tract and compute along-tract asymmetry profiles that quantify microstructural differences between hemispheres.

Studying asymmetry in Alzheimer’s disease (AD) is particularly important because AD pathology and clinical symptoms often emerge in a lateralized manner, with one hemisphere affected earlier or more severely than the other. Such hemispheric imbalances are closely linked to domain-specific deficits, including language, memory, and visuospatial function, which are each, to some extent, lateralized. Accordingly, numerous studies have examined gray- and white-matter asymmetry in AD using region-of-interest analyses or whole-tract average measures, demonstrating that disrupted hemispheric organization is a hallmark of the disease [8], [19]–[21].

More recently, tractometry has moved beyond whole-tract summaries toward along-tract profiling, enabling spatially localized characterization of microstructural degeneration along individual pathways [17], [22], [23]. However, these approaches typically analyze each tract independently within a single hemisphere and do not explicitly model hemispheric correspondence or asymmetry. As a result, conventional unilateral analyses can obscure hemispheric differences, potentially masking early or spatially focal disease effects. In this work, we address this gap by introducing an asymmetry-aware tractometry framework that explicitly models left–right correspondence along homologous pathways, providing a more sensitive and anatomically grounded tool for probing disease onset and progression in AD.

We apply our bundle asymmetry framework to diffusion MRI data from the Alzheimer’s Disease Neuroimaging Initiative (ADNI) to examine hemispheric imbalance in both mild cognitive impairment (MCI) and dementia, demonstrating how it reveals systematic left–right differences across major association, projection, and limbic pathways. Our method enables both subject-level and group-level quantification of asymmetry along the full extent of homologous tracts, providing a sensitive and anatomically grounded tool for studying white-matter asymmetry. This approach extends tractometry beyond unilateral analysis and opens new avenues to investigate hemispheric specialization and its disruption in neurological disorders.

## II. Methods

We analyzed 3D dMRI data from 1,215 participants, including 863 from ADNI-3 and 352 from ADNI-4, acquired using 10 distinct protocols across three scanner vendors: GE (General Electric), P (Philips), and S (Siemens). The protocols: GE127, GE36, GE54, P100, P33, P36, S100, S127, S31, and S55, represent different combinations of acquisition parameters, with the numeric suffix indicating the number of diffusion-weighted directions.

dMRI data were preprocessed using the ADNI dMRI protocol [24]–[26]. Preprocessing of raw dMRI data included denoising [27], [28], Gibbs deringing [29], [30], skull stripping [31], [32], motion and eddy current correction [32], [33], b1 bias field inhomogeneity correction [30], [34], and echo-planar imaging distortions correction. Diffusion tensor imaging (DTI) [35] models were fitted to estimate standard diffusion metrics, including fractional anisotropy (FA), mean diffusivity (MD), radial diffusivity (RD), and axial diffusivity (AxD).

### A. Bundle Atlas

To enable principled left–right comparisons of homologous white-matter bundles, we use a curated, symmetric version of the HCP1065 atlas [18], [36] that unifies the definitions of left and right tracts. In conventional atlases, hemispheric bundles are defined independently, which can introduce subtle geometric and topological discrepancies in bundle shape, extent, and orientation. Such differences can confound asymmetry analyses, as they may reflect atlas design rather than true biological variation.

**TABLE 1.**
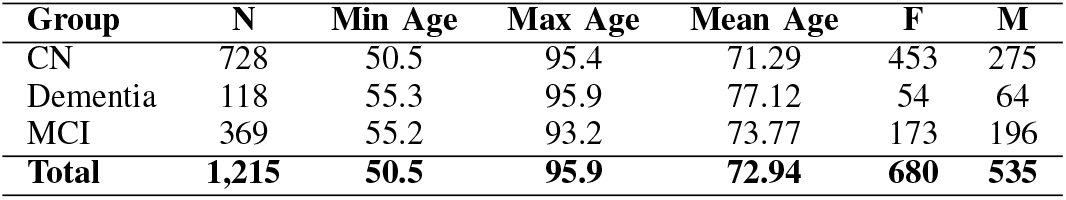
Demographic summary of the study cohort, reporting the distribution across diagnostic groups (Dx), age range, and sex, including counts of female (F) and male (M) participants.

We address this by adopting a symmetric atlas in which each homologous pair of bundles shares an identical geometric template in a common MNI space. Each hemispheric bundle is formed by combining the native tract with a mirrored version of its contralateral homolog. As illustrated in Figure 1 for the arcuate fasciculus, the left arcuate fasciculus (*AF*_*L*_) is obtained by joining the native *AF*_*L*_ with the mirrored *AF*_*R*_, while the right arcuate fasciculus (*AF*_*R*_) is created by combining the native *AF*_*R*_ with the mirrored *AF*_*L*_. This procedure ensures that both hemispheres share an identical geometric template while preserving their anatomical location. As a result, corresponding tracts in the left and right hemispheres have the same number of along-tract segments, an identical spatial extent, and a consistent parameterization along their length, as shown in the Figure. 2. Consequently, any measured hemispheric differences arise from subject-specific anatomy and microstructure rather than from atlas construction.

**Fig. 1:**
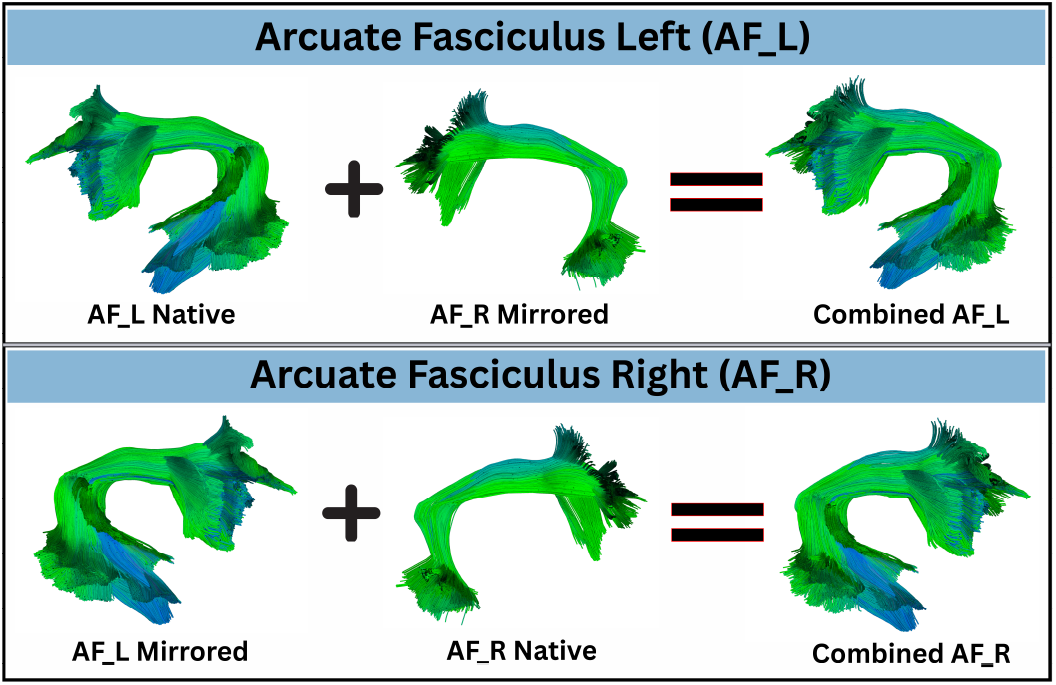
Symmetric Atlas Bundles: Illustration of the construction of a symmetric model for a single homologous pair using the arcuate fasciculus as an example. Each hemispheric bundle is formed by combining the native tract with a mirrored version of its contralateral homolog to enforce geometric symmetry. Specifically, the left arcuate fasciculus (*AF*_*L*_) is obtained by joining the native *AF*_*L*_ with the mirrored *AF*_*R*_ in a common symmetric space. Conversely, the right arcuate fasciculus (*AF*_*R*_) is created by combining the native *AF*_*R*_ with the mirrored *AF*_*L*_. This procedure ensures that structures in both hemispheres share an identical geometric template while preserving their anatomical location.

**Fig. 2:**
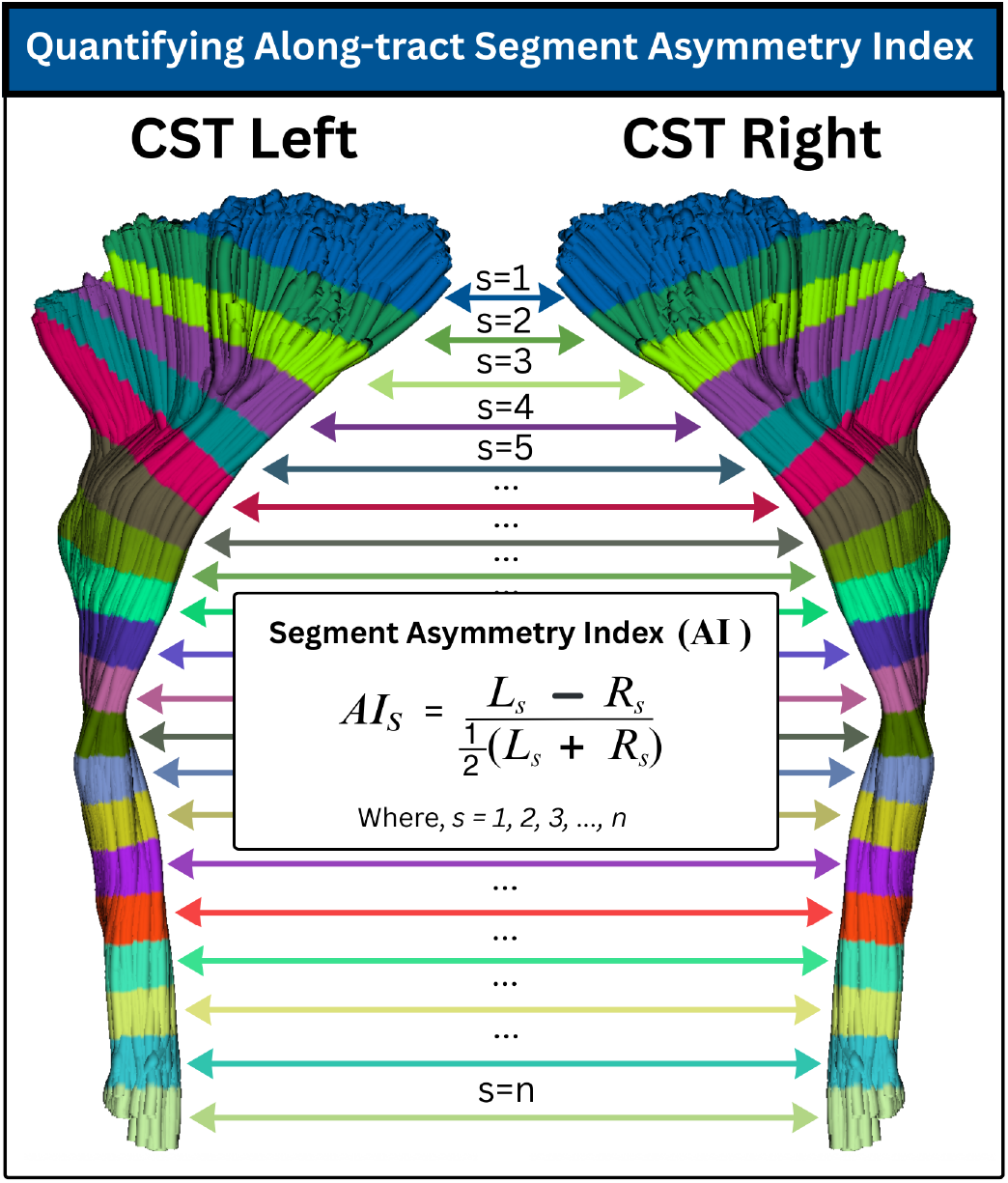
Bundle Segment-wise Asymmetry Index (AI) Illustration of the symmetric tractometry framework for computing along-tract hemispheric asymmetry. A homologous white-matter bundle (here, the corticospinal tract) is represented using a symmetric model, ensuring that left and right tracts share an identical geometric template and orientation. Each bundle is divided into *n* corresponding segments along its length, such that segment *s* in the left hemisphere maps to the same anatomical location as segment *s* in the right hemisphere. For each subject and each segment, microstructural values are extracted to form paired profiles *L*_*s*_ and *R*_*s*_. The segment-wise Asymmetry Index (AI) is then computed, yielding an anatomically aligned, along-tract measure of hemispheric asymmetry.

### B. Bundle Segmentation

We performed bundle-specific tractography and segmentation using a symmetric population atlas as an anatomical prior [37]. Spherical harmonics were computed using Constrained spherical deconvolution (CSD) [38], except for data acquired with multi-shell protocols (GE127, P100, S100, and S127), where Multi-Shell Multi-Tissue CSD [39] was used. For each subject, FA, MD, and mean b0 maps were nonlinearly registered to the ICBM 2009a MNI template using multichannel ANTs Syn [40]. Atlas bundles from the symmetric HCP-1065 atlas were converted into binary masks, dilated, and warped into each subject’s diffusion space using the inverse ANTs transform. These warped bundles defined bundle-specific seeding masks, stopping masks, and streamline length constraints. Within each seed mask, local probabilistic tractography was performed using DIPY [41] with bundle-specific angular thresholds (20^*°*^-30^*°*^) and a small step size (0.2mm). Streamlines were retained only if their lengths fell within the atlas-defined range with a tolerance margin. After tracking, the atlas bundles were warped into subject space and used to refine the tractograms using Fast Streamline Search [42] with custom distance thresholds (6-7mm), optimized per bundle. Note that the symmetric atlas bundles are used solely to *guide* tractography within anatomically plausible bundle-specific regions. They do not impose streamlines where none exist: if a pathway is absent or severely degraded in a subject, no streamlines are artificially constructed. Instead, the symmetric atlas provides each homologous tract in both hemispheres an equal opportunity to be reconstructed and segmented when streamlines are present. This design mitigates atlas-definition bias, as asymmetric bundle templates can preferentially favor one hemisphere over the other, potentially confounding true biological differences with atlas geometry.

In this paper, we focus on a set of eight bilateral white-matter pathways spanning major association, projection, and limbic systems: the Arcuate Fasciculus (AF), Inferior Fronto-Occipital Fasciculus (IFOF), Middle Longitudinal Fasciculus (MdLF), Uncinate Fasciculus (UF), and Extreme Capsule (EMC) as association tracts; the Corticospinal Tract (CST) and Optic Radiation (OR) as projection tracts; and the Fornix (FX) as a core limbic pathway.

### C. BUAN Tractometry and Bundle Asymmetry Quantification

We use BUAN tractometry to generate bundle profiles of subjects for each of the eight bilateral tracts. For each homologous bundle pair, bundle profiles are created by dividing each bundle into *n* segments along its length using the model bundle centroid. Crucially, we enforce a consistent orientation across hemispheres so that segment *s* = 1 in the left bundle corresponds to the same anatomical end of the tract as segment *s* = 1 in the right bundle. This orientation constraint guarantees one-to-one correspondence between homologous segments across hemispheres.

Bundle profiles were generated using four DTI–based microstructural measures: FA, MD, RD, and AxD. Although BUAN can incorporate measures derived from more advanced models, such as diffusion kurtosis imaging (DKI) [43] or neurite orientation dispersion and density imaging (NODDI) [44], we focus on these four well-established DTI metrics. Bundle profiles are constructed by dividing each bundle into *n* = 50 segments using the model bundle centroid defined in common space. We cluster the model bundle using QuickBundles [45] to obtain a centroid consisting of 50 points along the tract. For each subject, we compute Euclidean distances between every point on every streamline and the 50 centroid points. Each streamline point is assigned to the nearest centroid point, thereby receiving a segment index. Streamlines are not resampled, and their native point distributions are preserved; consequently, multiple points from a streamline may contribute to the same segment. Because segment assignment is performed in a common symmetric space, this procedure establishes consistent segment correspondence across subjects, groups, and hemispheres. Microstructural measures (e.g., FA) are then projected onto bundle points in native space and aggregated as weighted means within each segment, yielding a central along-tract profile with one value per segment [46].

For each subject and each bundle, this results in paired along-tract profiles for the two hemispheres,

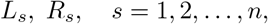

where *L*_*s*_ and *R*_*s*_ denote the microstructural value at segment *s* of the left and right bundle, respectively. Before computing hemispheric asymmetry, we apply tractometry-based ComBat harmonization [46], [47] to correct for scanner and protocol effects. Importantly, harmonization is performed on the left and right bundle profiles separately for each tract and metric. After these bundle-specific microstructural profiles are harmonized, we compute the segment-wise bundle asymmetry index, ensuring that measured hemispheric differences reflect biological variation rather than acquisition-related confounds.

### D. Along-Tract Bundle Asymmetry Index

We quantify hemispheric asymmetry in the microstructural measures (e.g., FA) at each segment using a normalized asymmetry index,

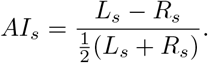

This formulation yields a scale-invariant measure of relative difference, where *AI*_*s*_ *>* 0 indicates leftward asymmetry and *AI*_*s*_ *<* 0 indicates rightward asymmetry at segment *s*. The result is an along-tract asymmetry profile,

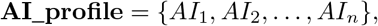

that captures spatially localized hemispheric differences along the full extent of each bundle. Figure. 2 provides a visual overview of the proposed along-tract asymmetry quantification.

These asymmetry profiles can be analyzed at the individual level or aggregated across subjects to study group-level patterns of hemispheric specialization and their alteration in aging and disease.

### E. Statistical Analysis

To assess group differences in along-tract hemispheric asymmetry, we use linear mixed-effects models (LMMs) independently at each along-tract segment.

For each bundle, metric, and segment *s* = 1, …, *n*, we model the asymmetry index *AI*_*i,s*_ of subject *i* as

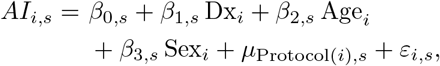

where Dx_*i*_ encodes diagnostic group (CN vs. MCI or CN vs. Dementia), Age_*i*_ and Sex_*i*_ are subject-level covariates, and 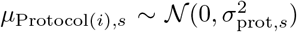 is a random intercept capturing scanner-protocol-specific effects. The residuals 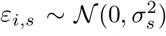 model unexplained variability.

The coefficient *β*_1,*s*_ tests for diagnosis-related changes in hemispheric asymmetry at segment *s*. We report the associated *p*-values and compute standardized effect sizes by normalizing *β*_1,*s*_ by the empirical standard deviation of *AI*_*·,s*_, enabling comparison across segments and bundles. To control for multiple comparisons along each tract, false discovery rate (FDR) correction [48] is applied across the *n* segment-wise tests per bundle and metric.

## III. Results and Discussion

Figure. 3 shows along-tract white-matter asymmetry for CN vs. Dementia, while Figure. 4 presents the corresponding results for CN vs. MCI. In each plot, the x-axis represents the along-tract segment index (*s* = 1, …, 50). The left y-axis reports the negative logarithm of the raw *p*-values, whereas the right y-axis shows the range of standardized effect sizes (standardized *β*). The red dotted horizontal line indicates the FDR–corrected significance threshold at *q* = 0.05. Blue bars depict *−*log_10_(*p*) for each segment, and the purple curve shows the standardized *β* coefficient, reflecting both the magnitude and direction of asymmetry between groups.

**Fig. 3:**
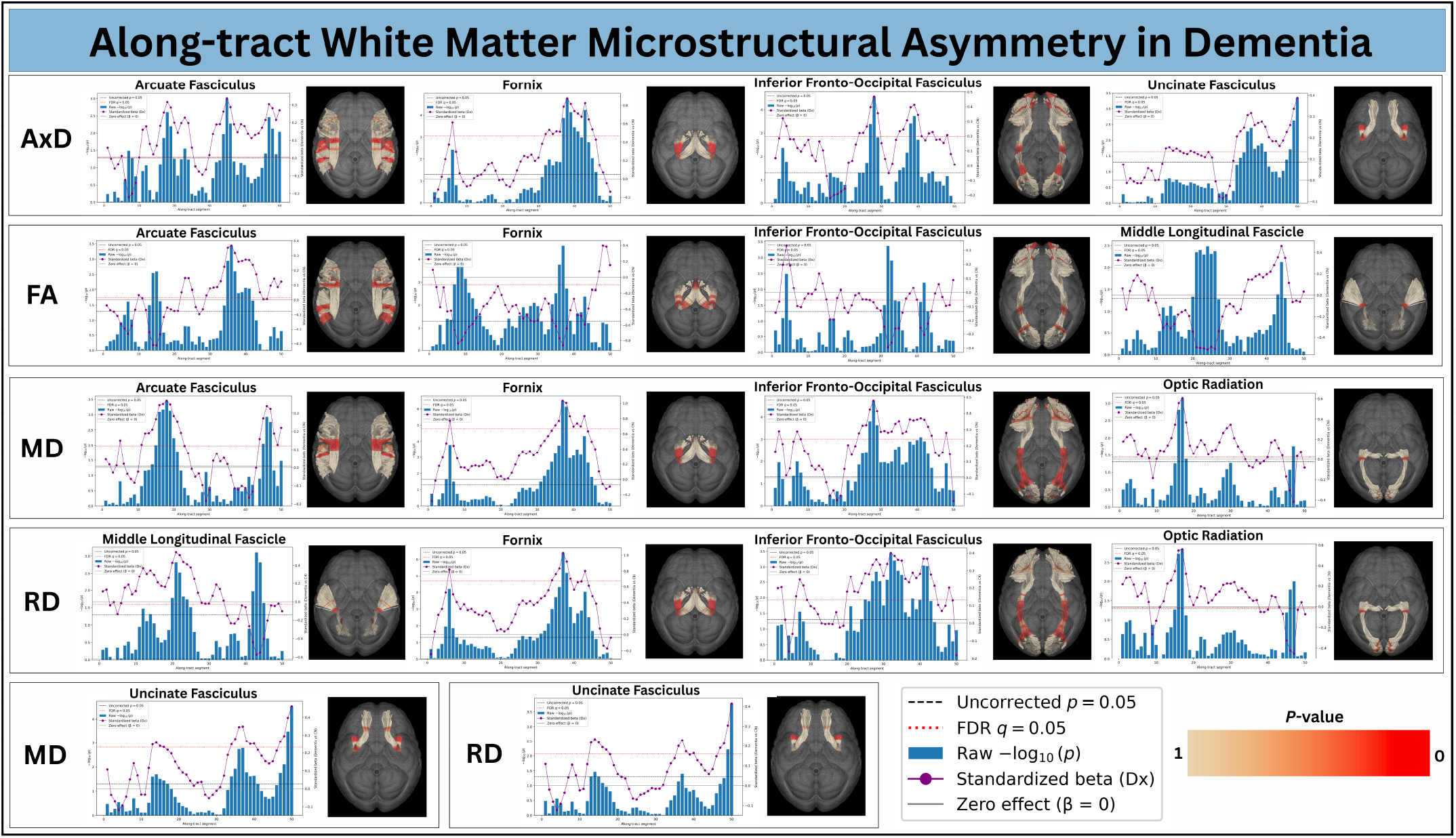
Along-tract Microstructural Asymmetry in Dementia. Segment-wise differences in left–right white-matter asymmetry between cognitively normal controls (CN) and individuals with dementia. For each bundle, the x-axis represents the along-tract segment index, and the left y-axis shows the negative logarithm of the raw *p*-values. Blue bars depict *−* log_10_(*p*) for each segment, with the dotted horizontal line marking the FDR–corrected significance threshold (*q* = 0.05). The right y-axis reports the standardized effect size, shown by the overlaid purple curve, which encodes both the magnitude and direction of asymmetry change. The accompanying three-dimensional renderings project the same segment-wise statistics onto homologous left and right tracts, highlighting the anatomical locations of significant effects in red color. Positive effect sizes indicate increased leftward asymmetry in dementia relative to controls, whereas negative values indicate increased rightward asymmetry.

**Fig. 4:**
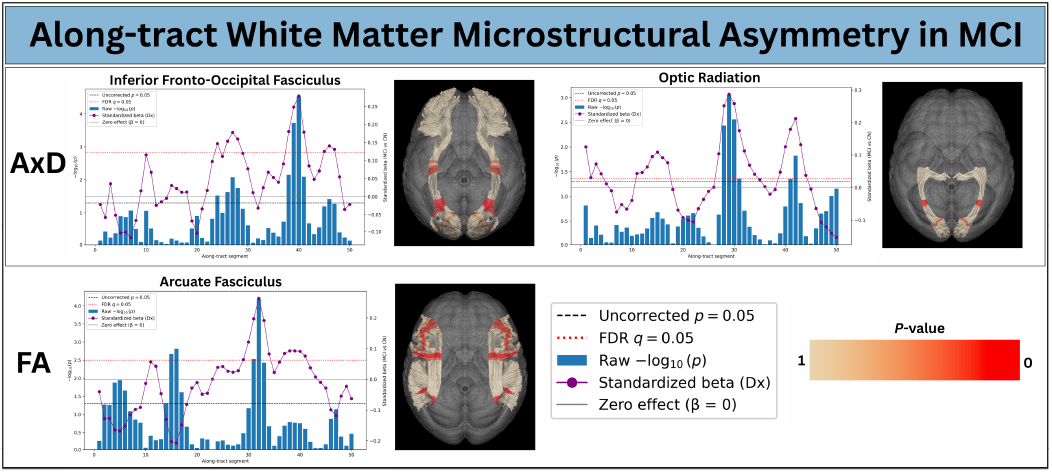
Along-tract Microstructural Asymmetry in Mild Cognitive Impairment (MCI). Segment-wise differences in left–right white-matter asymmetry between cognitively normal controls (CN) and individuals with MCI. For each bundle, the x-axis represents the along-tract segment index, and the left y-axis shows the negative logarithm of the raw *p*-values. Blue bars depict *−*log_10_(*p*) for each segment, with the dotted horizontal line marking the FDR–corrected significance threshold (*q* = 0.05). The right y-axis reports the standardized effect size, shown by the overlaid purple curve, which encodes both the magnitude and direction of asymmetry change. The accompanying three-dimensional renderings project the same segment-wise statistics onto homologous left and right tracts, highlighting the anatomical locations of significant effects in red color. Positive effect sizes indicate increased leftward asymmetry in MCI relative to controls, whereas negative values indicate increased rightward asymmetry.

Each along-tract plot is accompanied by a three-dimensional visualization of the corresponding white-matter bundles, in which the same *p*-values are projected onto both left and right hemispheric tracts. Red color on tracts indicates *p*-values *<* 0.05. This joint visualization highlights the specific along-tract regions exhibiting significant asymmetry, directly linking statistical effects to their anatomical locations.

In the CN vs. Dementia comparison, we observe widespread and spatially localized asymmetry across multiple major pathways, including the arcuate fasciculus, fornix, inferior fronto-occipital fasciculus, uncinate fasciculus, middle longitudinal fasciculus, and optic radiation. These effects are not uniform along each tract: significant segments cluster in anatomically meaningful subregions, demonstrating that hemispheric imbalance in dementia is highly focal rather than global. Across metrics, AxD and MD exhibit the strongest and most consistent asymmetry, while FA and RD show more localized but still robust effects. Several tract segments surpass both uncorrected and FDR-corrected thresholds, indicating pronounced disease-related divergence between homologous left and right pathways.

In contrast, the CN vs. MCI results reveal more selective and spatially restricted asymmetry. Significant effects are primarily confined to a smaller set of bundles, most notably the inferior fronto-occipital fasciculus, optic radiation, and arcuate fasciculus, and involve fewer contiguous segments than in Dementia. The magnitude of effects is reduced, consistent with MCI representing an intermediate stage between normal aging and dementia. Importantly, the same tracts that exhibit early asymmetry in MCI show broader and stronger effects in Dementia, suggesting a progressive expansion of hemispheric imbalance with disease severity. However, it is important to note that the MCI and Dementia cohorts consist of different individuals; the two groups do not represent longitudinal progression within the same subjects, but rather cross-sectional comparisons between distinct populations.

The standardized beta curves provide the direction of asymmetry change. Positive betas indicate a shift toward relatively greater left-hemisphere values compared to controls, whereas negative betas reflect a relative right-hemisphere dominance. The coexistence of positive and negative regions along the same tract demonstrates that disease-related asymmetry is not a simple global left–right shift, but a complex, segment-specific reorganization in which different subregions diverge in opposite directions. These directional changes align with peaks in statistical significance, reinforcing that the observed effects reflect systematic, anatomically localized alterations rather than noise.

Long association pathways such as the arcuate fasciculus, inferior fronto-occipital fasciculus, uncinate fasciculus, and middle longitudinal fasciculus exhibit alternating left-heavy and right-heavy segments along their trajectories. Across the four DTI metrics, most regions in these tracts show positive *β* values, indicating a leftward shift in hemispheric balance in Dementia relative to controls, while select segments shift toward negative *β*, reflecting rightward shifts. This pattern implies that fronto–temporal and fronto–parietal systems supporting language, memory, and higher-order cognition, domains that are themselves lateralized, are disrupted in a spatially heterogeneous manner [49]–[51].

Asymmetry in the fornix is dominated by left-heavy significant segments in MD and RD, accompanied by right-heavy effects in FA, pointing to differential hemispheric involvement of limbic memory circuitry [52]. In contrast, the optic radiation is predominantly left-heavy, with positive *β* peaks across most significant segments in MD and RD, yet also exhibits focal right-heavy subregions. Together, these patterns indicate that even primary projection systems [53] exhibit hemispheric imbalance, and that disease-related asymmetry is not uniform along a tract but varies across its subcomponents.

Together, these findings show that neurodegenerative disease is associated with structured, along-tract microstructural asymmetry that emerges in MCI and becomes more extensive in Dementia. By combining symmetric tract definitions, tract-specific harmonization, and along-tract mixed-effects modeling, our framework reveals where, and in which direction, hemispheric balance is disrupted, providing a fine-grained map of disease-related white-matter reorganization.

Future work will extend this framework to a broader repertoire of white-matter pathways, including commissural and cerebellar tracts, to provide a more comprehensive map of hemispheric imbalance. In addition to microstructural metrics, we will integrate geometric [54] and morphometric [55] features, such as tract shape, curvature, and bundle morphometry profile, into the asymmetry analysis. Incorporating these complementary dimensions will enable joint modeling of microstructural and structural divergence along homologous pathways, yielding a richer and more mechanistic characterization of how neurodegenerative processes reshape white-matter organization across hemispheres.

## Conclusion

We introduced a tractometry framework for quantifying along-tract hemispheric asymmetry in white matter that is principled, anatomically consistent, and robust to multi-site acquisition effects. By leveraging a BUAN tractometry, symmetric population-based atlas, enforcing segment-wise correspondence across hemispheres, and harmonizing tract profiles prior to asymmetry computation, our approach ensures that left–right differences reflect subject-specific neurobiology rather than atlas design or scanner variability. This enables fine-grained, anatomically meaningful comparisons of homologous white-matter fiber bundles along their full extent.

Applying this bundle asymmetry framework to ADNI dMRI data revealed spatially localized patterns of microstructural asymmetry that differentiate cognitively normal controls from individuals with MCI and dementia. These effects are not uniform along tracts, underscoring the importance of along-tract analysis over region-averaged or ROI-based asymmetry measures.

Our results demonstrate that white-matter asymmetry is a sensitive marker of neurodegenerative change, even at prodromal stages, and that its spatial organization carries biologically meaningful information. More broadly, this work establishes a general framework that can be applied across cohorts, scanners, and clinical populations. Integrating bundle asymmetry quantification into BUAN tractometry provides a scalable investigation of hemispheric specialization and its disruption in brain disease. As such, it might open up new avenues for discovering spatially specific biomarkers of neurological and psychiatric disorders.

## Acknowledgment

This research was supported by the NIH (National Institutes of Health) under grants U01 AG068057, RF1 NS136995, P41 EB015922, RF1 AG057892, R01 MH134004, and R01 EB027585, and the US Alzheimer’s Association under grant AARG-23-1149996.

## References

[1] A. W. Toga and P. M. Thompson, “Mapping brain asymmetry,” Nature Reviews Neuroscience, vol. 4, no. 1, pp. 37–48, 2003.

[2] M. C. Corballis, “The evolution and genetics of cerebral asymmetry,” Philosophical Transactions of the Royal Society B: Biological Sciences, vol. 364, no. 1519, pp. 867–879, 2009.

[3] M. Annett, Handedness and brain asymmetry: The right shift theory. Psychology Press, 2013.

[4] A. Bethmann, C. Tempelmann, R. De Bleser, H. Scheich, and A. Brechmann, “Determining language laterality by fmri and dichotic listening,” Brain Research, vol. 1133, pp. 145–157, 2007.

[5] I. Andrulyte, E. Demirkan, F. M. Branzi, L. J. Bonnett, and S. S. Keller, “Does white matter structure relate to hemispheric language lateralisation? a systematic review,” Brain Communications, p. fcag009, 01 2026.

[6] L. Dorfschmidt, S. White, M. Gardner, S. Bedford, G. Ball, A. D. Edwards, Y. Gu, Y. He, H. Huang, S. Karandikar, et al., “Charting structural brain asymmetry across the human lifespan,” bioRxiv, 2025.

[7] P. Kanakaraj, S. Bogdanov, M. E. Kim, J. Samir, C. Gao, K. Ramadass, G. Rudravaram, N. R. Newlin, D. Archer, T. J. Hohman, et al., “Lifespan trajectories of asymmetry in white matter tracts,” bioRxiv, pp. 2025–09, 2025.

[8] P. M. Thompson, K. M. Hayashi, G. De Zubicaray, A. L. Janke, S. E. Rose, J. Semple, D. Herman, M. S. Hong, S. S. Dittmer, D. M. Doddrell, et al., “Dynamics of gray matter loss in alzheimer’s disease,” Journal of Neuroscience, vol. 23, no. 3, pp. 994–1005, 2003.

[9] Y. Liu, J. Yuan, C. Tan, M. Wang, F. Zhou, C. Song, Y. Tang, X. Li, Q. Liu, Q. Shen, et al., “Exploring brain asymmetry in early-stage parkinson’s disease through functional and structural mri,” CNS Neuroscience & Therapeutics, vol. 30, no. 7, p. e14874, 2024.

[10] M. C. Postema, D. Van Rooij, E. Anagnostou, C. Arango, G. Auzias, M. Behrmann, G. B. Filho, S. Calderoni, R. Calvo, E. Daly, et al., “Altered structural brain asymmetry in autism spectrum disorder in a study of 54 datasets,” Nature Communications, vol. 10, no. 1, p. 4958, 2019.

[11] C. Banfi, K. Koschutnig, K. Moll, G. Schulte-Körne, A. Fink, and K. Landerl, “White matter alterations and tract lateralization in children with dyslexia and isolated spelling deficits,” Human Brain Mapping, vol. 40, no. 3, pp. 765–776, 2019.

[12] A. Yendiki, P. Panneck, P. Srinivasan, A. Stevens, L. Zöllei, J. Augustinack, R. Wang, D. Salat, S. Ehrlich, T. Behrens, et al., “Automated probabilistic reconstruction of white-matter pathways in health and disease using an atlas of the underlying anatomy,” Frontiers in neuroinformatics, vol. 5, p. 23, 2011.

[13] J. D. Yeatman, R. F. Dougherty, N. J. Myall, B. A. Wandell, and H. M. Feldman, “Tract profiles of white matter properties: automating fiber-tract quantification,” PloS one, vol. 7, no. 11, p. e49790, 2012.

[14] B. Q. Chandio, S. L. Risacher, F. Pestilli, D. Bullock, F.-C. Yeh, S. Koudoro, A. Rokem, J. Harezlak, and E. Garyfallidis, “Bundle analytics, a computational framework for investigating the shapes and profiles of brain pathways across populations,” Scientific Reports, vol. 10, no. 1, p. 17149, 2020.

[15] G. S. Kim, B. Q. Chandio, S. M. Benavidez, Y. Feng, P. M. Thompson, and K. E. Lawrence, “Mapping along-tract commissural and association white matter microstructural differences in autistic children and young adults,” Cerebral Cortex, vol. 35, no. 11, p. bhaf291, 2025.

[16] K. Chang, L. Burke, N. LaPiana, B. Howlett, D. Hunt, M. Dezelar, J. B. Andre, P. Curl, J. D. Ralston, A. Rokem, et al., “Free water elimination tractometry for aging brains,” Imaging Neuroscience, vol. 3, pp. IMAG–a, 2025.

[17] B. Q. Chandio, C. Owens-Walton, J. E. Villalon-Reina, L. Nabulsi, S. I. Thomopoulos, J. Guaje, E. Garyfallidis, and P. M. Thompson, “Microstructural changes in the white matter tracts of the brain due to mild cognitive impairment,” Alzheimer’s & Dementia, vol. 18, p. e065339, 2022.

[18] F.-C. Yeh, “Population-based tract-to-region connectome of the human brain and its hierarchical topology,” Nature Communications, vol. 13, no. 1, p. 4933, 2022.

[19] P. Thompson, J. Moussai, S. Zohoori, A. Goldkorn, A. Khan, M. Mega, G. Small, J. Cummings, and A. Toga, “Cortical variability and asymmetry in normal aging and alzheimer’s disease.,” Cerebral Cortex (New York, NY: 1991), vol. 8, no. 6, pp. 492–509, 1998.

[20] S. Derflinger, C. Sorg, C. Gaser, N. Myers, M. Arsic, A. Kurz, C. Zimmer, A. Wohlschläger, and M. Mühlau, “Grey-matter atrophy in alzheimer’s disease is asymmetric but not lateralized,” Journal of Alzheimer’s Disease, vol. 25, no. 2, pp. 347–357, 2011.

[21] C. Wachinger, D. H. Salat, M. Weiner, M. Reuter, and A. D. N. Initiative, “Whole-brain analysis reveals increased neuroanatomical asymmetries in dementia for hippocampus and amygdala,” Brain, vol. 139, no. 12, pp. 3253–3266, 2016.

[22] B. Q. Chandio, J. E. Villalon-Reina, T. M. Nir, S. I. Thomopoulos, Y. Feng, S. Benavidez, N. Jahanshad, J. Harezlak, E. Garyfallidis, P. M. Thompson, et al., “Amyloid, tau, and apoe in alzheimer’s disease: Impact on white matter tracts,” Pacific Symposium on Biocomputing (PSB), 2025.

[23] B. Q. Chandio, T. M. Nir, J. E. Villalon-Reina, S. I. Thomopoulos, Y. Feng, R. I. Reid, C. R. Jack, M. W. Weiner, E. Garyfallidis, N. Jahanshad, et al., “White matter tract vulnerability to amyloid pathology on the alzheimer’s disease continuum,” in 2025 21st International Symposium on Biomedical Image Processing and Analysis (SIPAIM), pp. 1–5, IEEE, 2025.

[24] N. Jahanshad, P. V. Kochunov, E. Sprooten, R. C. Mandl, T. E. Nichols, L. Almasy, J. Blangero, R. M. Brouwer, J. E. Curran, G. I. de Zubicaray, et al., “Multi-site genetic analysis of diffusion images and voxelwise heritability analysis: A pilot project of the ENIGMA–DTI working group,” NeuroImage, vol. 81, pp. 455–469, 2013.

[25] S. I. Thomopoulos, T. M. Nir, J. E. V. Reina, N. Jahanshad, and P. M. Thompson, “Diffusion MRI metrics of brain microstructure in alzheimer’s disease: Boosting disease sensitivity with multi-shell imaging and advanced pre-processing: Neuroimaging/new imaging methods,” Alzheimer’s & Dementia, vol. 16, p. e046654, 2020.

[26] K. Liou, S. I. Thomopoulos, H. Yoo, Y. Shuai, S. Chehrzadeh, A. Arani, B. Borowski, R. I. Reid, C. R. Jack, M. W. Weiner, N. Jahanshad, P. M. Thompson, and T. M. Nir, “Dti versus noddi white matter microstructural biomarkers of alzheimer’s disease,” in 2025 21st International Symposium on Biomedical Image Processing and Analysis (SIPAIM), pp. 1–4, 2025.

[27] J. V. Manjón, P. Coupé, L. Concha, A. Buades, D. L. Collins, and M. Robles, “Diffusion weighted image denoising using overcomplete local PCA,” PLoS ONE, vol. 8, no. 9, 2013.

[28] E. Garyfallidis, M. Brett, B. Amirbekian, A. Rokem, S. Van Der Walt, M. Descoteaux, I. Nimmo-Smith, and D. Contributors, “Dipy, a library for the analysis of diffusion MRI data,” Frontiers in Neuroinformatics, vol. 8, p. 8, 2014.

[29] E. Kellner, B. Dhital, V. G. Kiselev, and M. Reisert, “Gibbs-ringing artifact removal based on local subvoxel-shifts,” Magnetic Resonance in Medicine, vol. 76, no. 5, pp. 1574–1581, 2016.

[30] J.-D. Tournier, R. Smith, D. Raffelt, R. Tabbara, T. Dhollander, M. Pietsch, D. Christiaens, B. Jeurissen, C.-H. Yeh, and A. Connelly, “MRtrix3: a fast, flexible and open software framework for medical image processing and visualisation,” NeuroImage, vol. 202, p. 116137, 2019.

[31] S. M. Smith, “Fast robust automated brain extraction,” Human Brain Mapping, vol. 17, no. 3, pp. 143–155, 2002.

[32] M. Jenkinson, C. F. Beckmann, T. E. Behrens, M. W. Woolrich, and S. M. Smith, “FSL,” NeuroImage, vol. 62, no. 2, pp. 782–790, 2012.

[33] J. L. Andersson and S. N. Sotiropoulos, “An integrated approach to correction for off-resonance effects and subject movement in diffusion MR imaging,” NeuroImage, vol. 125, pp. 1063–1078, 2016.

[34] N. J. Tustison, B. B. Avants, P. A. Cook, Y. Zheng, A. Egan, P. A. Yushkevich, and J. C. Gee, “N4itk: improved N3 bias correction,” IEEE Transactions on Medical Imaging, vol. 29, no. 6, pp. 1310–1320, 2010.

[35] P. J. Basser, J. Mattiello, and D. LeBihan, “MR diffusion tensor spectroscopy and imaging,” Biophysical Journal, vol. 66, no. 1, pp. 259–267, 1994.

[36] I. Ba Gari, J. Davis Alibrando, Y. Feng, J. E. Villalon-Reina, A. Cabalinan, L. Nabulsi, P. M. Thompson, F.-C. Yeh, B. Q. Chandio, T. M. Nir, and N. Jahanshad, “Enigma symmetric white matter tractography atlas,” Apr. 2026.

[37] Y. Feng, Y. Shuai, J. E. Villalón-Reina, B. Q. Chandio, S. I. Thomopoulos, T. M. Nir, N. Jahanshad, and P. M. Thompson, “Streamline density normalization: A robust approach to mitigate bundle variability in multi-site diffusion mri,” in International Workshop on Computational Diffusion MRI, pp. 44–56, Springer, 2025.

[38] J.-D. Tournier, F. Calamante, and A. Connelly, “Robust determination of the fibre orientation distribution in diffusion mri: non-negativity constrained super-resolved spherical deconvolution,” NeuroImage, vol. 35, no. 4, pp. 1459–1472, 2007.

[39] B. Jeurissen, J.-D. Tournier, T. Dhollander, A. Connelly, and J. Sijbers, “Multi-tissue constrained spherical deconvolution for improved analysis of multi-shell diffusion MRI data,” NeuroImage, vol. 103, pp. 411–426, 2014.

[40] B. B. Avants, N. Tustison, G. Song, et al., “Advanced normalization tools (ants),” Insight j, vol. 2, no. 365, pp. 1–35, 2009.

[41] E. Garyfallidis, M. Brett, B. Amirbekian, A. Rokem, S. van der Walt, M. Descoteaux, I. Nimmo-Smith, and Dipy Contributors, “Dipy, a library for the analysis of diffusion MRI data,” Frontiers in Neuroinformatics, vol. 8, Feb. 2014.

[42] E. St-Onge, E. Garyfallidis, and D. L. Collins, “Fast streamline search: an exact technique for diffusion mri tractography,” Neuroinformatics, vol. 20, no. 4, p. 1093, 2022.

[43] A. J. Steven, J. Zhuo, and E. R. Melhem, “Diffusion kurtosis imaging: an emerging technique for evaluating the microstructural environment of the brain,” American Journal of Roentgenology, vol. 202, no. 1, pp. W26–W33, 2014.

[44] H. Zhang, T. Schneider, C. A. Wheeler-Kingshott, and D. C. Alexander, “NODDI: practical in vivo neurite orientation dispersion and density imaging of the human brain,” NeuroImage, vol. 61, no. 4, pp. 1000–1016, 2012.

[45] E. Garyfallidis, M. Brett, M. M. Correia, G. B. Williams, and I. Nimmo-Smith, “Quickbundles, a method for tractography simplification,” Frontiers in Neuroscience, vol. 6, p. 175, 2012.

[46] B. Q. Chandio, J. E. Villalon-Reina, T. M. Nir, S. I. Thomopoulos, Y. Feng, S. Benavidez, N. Jahanshad, J. Harezlak, E. Garyfallidis, and P. M. Thompson, “Bundle analytics based data harmonization for multi-site diffusion mri tractometry,” in 2024 46th Annual International Conference of the IEEE Engineering in Medicine and Biology Society (EMBC), pp. 1–7, IEEE, 2024.

[47] J.-P. Fortin, N. Cullen, Y. I. Sheline, W. D. Taylor, I. Aselcioglu, P. A. Cook, P. Adams, C. Cooper, M. Fava, P. J. McGrath, et al., “Harmonization of cortical thickness measurements across scanners and sites,” NeuroImage, vol. 167, pp. 104–120, 2018.

[48] Y. Benjamini and Y. Hochberg, “Controlling the false discovery rate: a practical and powerful approach to multiple testing,” Journal of the Royal Statistical Society: series B (Methodological), vol. 57, no. 1, pp. 289–300, 1995.

[49] M. Catani and M. T. De Schotten, “A diffusion tensor imaging tractography atlas for virtual in vivo dissections,” Cortex, vol. 44, no. 8, pp. 1105–1132, 2008.

[50] H. Duffau, “Stimulation mapping of white matter tracts to study brain functional connectivity,” Nature Reviews Neurology, vol. 11, no. 5, pp. 255–265, 2015.

[51] F. Latini, G. Trevisi, M. Fahlström, M. Jemstedt, Å. Alberius Munkhammar, M. Zetterling, G. Hesselager, and M. Ryttlefors, “New insights into the anatomy, connectivity and clinical implications of the middle longitudinal fasciculus,” Frontiers in Neuroanatomy, vol. 14, p. 610324, 2021.

[52] J. P. Aggleton, S. M. O’Mara, S. D. Vann, N. F. Wright, M. Tsanov, and J. T. Erichsen, “Hippocampal–anterior thalamic pathways for memory: uncovering a network of direct and indirect actions,” European Journal of Neuroscience, vol. 31, no. 12, pp. 2292–2307, 2010.

[53] A. J. Sherbondy, R. F. Dougherty, S. Napel, and B. A. Wandell, “Identifying the human optic radiation using diffusion imaging and fiber tractography,” Journal of vision, vol. 8, no. 10, pp. 12–12, 2008.

[54] F.-C. Yeh, “Shape analysis of the human association pathways,” NeuroImage, vol. 223, p. 117329, 2020.

[55] B. Q. Chandio, E. Olivetti, D. Romero-Bascones, S. I. Thomopoulos, J. E. Villalon-Reina, T. M. Nir, J. Harezlak, P. M. Thompson, and E. Garyfallidis, “Bundlewarp: Enhancing white matter tractometry and morphometry with precise neuronal mapping using streamline-based nonlinear registration,” Medical Image Analysis, p. 104114, 2026.

